# MiR-135-5p alleviates bone cancer pain by regulating astrocyte-mediated neuroinflammation in spinal cord through JAK2/STAT3 signaling pathway

**DOI:** 10.1101/2021.05.17.444477

**Authors:** Ming Liu, Xuefeng Cheng, Hong Yan, Jingli Chen, Caihua Liu, Zhonghui Chen

**Affiliations:** Department of Anaesthesiology, The Central Hospital of Wuhan, Tongji Medical College, Huazhong University of Science and Technology; Department of Spine Surgery, Renmin Hospital of Wuhan University

**Keywords:** miRNA-135-5P, JAK2/STAT3 signaling pathway, neuroinflammation, astrocyte activation, bone cancer pain, mouse model

## Abstract

Bone cancer pain (BCP) was associated with microRNA dysregulation. In this study, we intended to clarify the potential role of miR-135-5p in a BCP mouse model, which was established by tumor cell implantation (TCI) in the medullary cavity of the mouse femur. The BCP related behaviors were tested, including the paw withdrawal mechanical threshold (PWMT) and number of spontaneous flinches (NSF). The miRNA expression profiles in astrocytes of the sham and tumor groups were compared, and miRNA microarray and quantitative real-time PCR (qRT-PCR) assays confirmed that the amount of expression of miR-135-5p was significantly decreased in astrocytes of the tumor group. Gain- and loss-of-function studies showed that miR-135-5p could inhibit astrocytes activation and inflammation cytokines (TNF-α and IL-1β) expression. The relation between miR-135-5p and JAK2 was detected by bioinformatic analysis and dual luciferase reporter gene assay. By conducting in vitro experiments, it was shown that the miR-135-5P mimics lowered the level of JAK2/STAT3 proteins and inflammatory factors in astrocytes. Moreover, in vivo analysis on BCP mice model indicated that the miR-135-5p agonist could sufficiently increase PWMT and decrease NSF. Meanwhile, reduced activation of astrocytes in the spinal cord, as well as decreased expression of JAK2/STAT3 and inflammatory mediators, were found after miR-135-5p agonist treatment. Collectively, the results showed that miR-135-5p could potentially reduce BCP in mice through inhibiting astrocyte-mediated neuroinflammation and blocking of the JAK2/STAT3 signaling pathway, indicating that the upregulation of miR-135-5P could be a therapeutic focus in BCP treatment.

## Introduction

Bone cancer pain (BCP), a complication that is encountered most often when metastasis of tumors to the bone occur, can lead to anxiety, depression, and other types of disorders in patients and could even greatly weaken their quality of life and functional status [1,2]. Since limitations exist in the current available treatment, many BCP patients have to endure limited relief of their pain and negative side effects. Therefore, knowledge of the potential molecular and cellular mechanisms of BCP is needed to be able to effectively treat these patients.

The pathogenesis of BCP is related to a multitude of genetical and biological regulators [3,4], however, recently an increasing amount of evidence support that microRNAs (miRNAs) play an essential role as regulators in BCP development [5-8]. MiRNAs can regulate multiple pathological and physiological processes, including differentiation, morphogenesis, apoptosis, proliferation, and survival on the protein level, through binding to the 3’ untranslated region of their target mRNAs [9]. Previous research has verified that miR-135-5p is downregulated in spinal cord tissue that has been injured, stimulates regrowth of axon, and has a neuroprotective role due to the inhibition of cellular apoptosis [10,11]. However, to date there is not one study that has investigated the mechanisms of miR-135-5p in BCP, particularly in relation to the signaling pathway and its target genes.

Previous research has shown that astrocytes have a key role in multiple biological functions, including modulation and perception of pain [12-14]. Microglia and astrocytes in the spinal cord also have a role to play in triggering and maintaining continuous pain originating from nerve injury and inflamed tissue [15-17]. It was discovered that in multiple pathological pain conditions astrocytes were activated, which was attributed to hypertrophy and the upregulation of glial fibrillary acidic protein (GFAP), and commonly viewed as the cause of intensifying persistent states of pain [18]. Moreover, astrocytes that are activated can discharge different inflammatory cytokines, like IL-1β and TNF-α[19,4]. The regulation of pain sensitivity is related to the assembly of proinflammatory mediators, resulting in central and peripheral sensitization, and the initiation of chronic pain [20].

Many studies have investigated which roles various intracellular transduction pathways and extracellular signaling molecules play in chronic pain conditions [21,4,22,23]. The Janus tyrosine kinase (JAK) family of activators and signal transducers of the transcription (STAT) pathway are considered as one of the most significant pathways of cell signaling in chronic pain conditions, which involves 7 different types of STAT and 4 different types of JAK [24,14]. A previous study revealed that the astrocytic JAK-STAT3 signaling pathway, in rats, can regulate proliferation of the spinal astrocyte and maintenance of neuropathic pain [14]. JAK2/STAT3 has demonstrated its role as a potential target in neuropathic pain therapy [25-27].

According to the forementioned, we hypothesized that miR-135-5p could affect BCP development involving the activation of astrocytes and the astrocytic JAK2/STAT3 signaling pathway. Therefore, the objective of this study is to evaluate through which mechanism the miR-135-5p/JAK2/STAT3 axis is involved in regulating BCP.

## Materials and methods

### Animals

The experimental protocols have been authorized by the Institutional Animal Care and Use Committee of our hospital and performed according to the guidelines of the International Association for the Study of Pain. C3H/HeJ mice, all male, (of 5□weeks old and 20□25 g) were acquired from the Beijing Vital River Laboratory Animal Technology Co., Ltd., (Beijing, China). The mice were kept inside a room which had temperature control (21±1°C) and alternating 12 h dark/light sequences. Everything possible was undertaken to reduce both animal suffering and the number of mice used to a minimum.

### BCP model

NCTC 135 medium (Sigma, USA) supplemented with 10% of horse serum (Gibco, USA) was used to culture the NCTC 2472 osteolytic sarcoma cells (American Type Culture Collection, USA), which were placed in an atmosphere of 37°C (Thermo Forma, USA) with 5% CO_2_. The steps to produce a BCP model have been previously described by Schwei et al [28]. First, anesthesia in the mice was achieved by intraperitoneally injecting them with a dose of 50 mg/kg pentobarbital sodium. Following general anesthesia, arthrotomy of the right knee was conducted. Next, we injected a dose of 20 μl α–minimum essential medium (Thermo Fisher Scientific, USA), which contained 2×10^5^ osteolytic sarcoma cells, inside the right femur’s intramedullary space, while the sham group was injected with just a dose of 20 μl α–minimum essential medium. Following the use of bone wax to seal off the drill hole, wound closure was achieved using 4-0 silk sutures (Ethicon, USA). The mice were transferred to a heated blanket to recover from anesthesia.

### Drug administration

To perform the therapeutic experiments, 48 male mice were selected at random and separated into four groups (n = 12 per group) to undergo sham or tumor cell implantation (TCI) surgery. Blinded observers performed the experiments as follows: (1) sham+vehicle group (the mice received sham surgery and were treated with 5 μl of PBS), (2) sham+agomiR NC group (the mice received sham surgery and were treated with 5 μl of agomiR NC [RiboBio, Guangzhou, China]), (3) tumor+agomiR NC group (the mice received TCI surgery and were treated with 5 μl of agomiR NC), and (4) tumor+agomiR-135-5p group (the mice received TCI surgery and were treated with 5 μl of agomiR-135-5p [RiboBio, Guangzhou, China]). The intrathecal injection was performed through a lumbar puncture in the L4□5 or L5□6 intervertebral space every 2 days (three injections).

### Assessing bone cancer-related pain behaviors

Spontaneous pain and mechanical allodynia in the mice were assessed pre-surgery (on day 0) in addition to 4, 7, 10, 14, 21, and 28 days after surgery in each group, and 0, 3, 5, 7, 10, and 15 h after administration of agomiR-135-5p, agomiR NC, and vehicle. The experimenters were all blinded to the data regarding details of group assignment during behavioral tests.

#### Paw withdrawal mechanical threshold (PWMT)

PWMTs of the hind paw on the right side were assessed with von Frey filaments (0.16, 0.4, 0.6, 1.0, 1.4, and 2.0 g; Stoelting, USA) and the up-and-down method. The mice were first transferred to transparent plexiglass compartments containing a wire mesh floor to undergo an acclimatization period of 30 min, then the von Frey filaments were placed vertically to their plantar surface, followed by the recording of the lowest strength of filament stimulus that resulted in withdrawal or flinching of the paw.

#### Number of spontaneous flinches (NSF)

The mice were first transferred to clear plexiglass compartments containing a wire mesh floor to undergo an acclimatization period of 30 min, followed by assessment of the NSF of the hind paw on the right side for more than 2 min. All the mice were tested five times.

### Histological observation

Histological observation of the bones containing tumor was conducted on the 14th day following TCI surgery. The mice were first anesthetized and then perfused transcardially with 0.9% saline and subsequently 4% paraformaldehyde. Following removal of the tibia, it was first demineralized in ethylene diamine tetraacetic acid (EDTA) (10%) for the duration of 1 week and then fixed in paraffin. A microtome was first used to cut sections of 5-μm, which were stained with hematoxylin and eosin and observed under a microscope.

### Primary astrocyte culture and transfection

Spinal cord segments at L4 to L5 of the mice were removed and dissected. Following enzymatic digestion, the detached cells were placed in Dulbecco’s modified Eagle’s medium (DMEM), which had fetal bovine serum (10%), streptomycin (0.1 mg/mL), and penicillin (200 IU) added to it. Then, the cells were placed in 75 cm^2^ containers and subsequently cultured in a humidified incubator for 10-14 days at 37°C (5% CO^2^ /95% air). The containers were shaken, after 6 days, at a frequency of 220 rpm (for 24 h at 37°C) to yield purified astrocytes. Production of astrocyte-defined medium was achieved according to the following formulation: DMEM with fibronectin (1.5 μg/mL), transferrin (50 μg/mL), heparin sulfate (0.5 g/mL), sodium selenite (5.2 ng/mL), insulin (5 μg/mL), epidermal growth factor (10 ng/mL), and basic fibroblast growth factor (5 ng/mL) added to it. The medium was changed after the first 3 days and three times a week thereafter. Lipofectamine 3000 (Invitrogen, Carlsbad, CA, USA) was utilized to transfect the astrocytes with miR-135-5p mimics, a mimic NC, a miR-135-5p inhibitor, and an inhibitor NC (RiboBio, Guangzhou, China), at a total concentration of 100 nM for 6 h.

### RNA isolation, cDNA synthesis, and qRT-PCR

Trizol reagent (Invitrogen, USA) was utilized to obtain total RNA from the spinal cords of the mice. The One-step PrimeScript miRNA cDNA Synthesis Kit (Takara, Japan) was utilized for reverse transcription. In accordance with the primer set dedicated to the target mRNA, iQSYBR Green mix (Bio-Rad, USA) was used to facilitate amplification of fragments by real-time quantitative reverse transcription PCR (RT-qPCR). A PCR flatform (Applied Biosystems, USA), capable of taking measurements simultaneously, was used to carry out the reactions. The quantitative results in raw form were first standardized to U6 level, followed by processing of the data with the Ct (ΔΔCt) method (2-ΔΔCt including logarithmic transformation) in order to approximate the relative expression levels.

### Microarray analyses

Firstly, Trizol method was used for extracting of total RNA from the L4 to L5 spinal cord segments with a total concentration of 1 mg/ml. Then, to purify the miRNA part of the total RNA a miRNA isolation kit (Ambion, USA) of mir Vana TM was used. To analyze the extracted miRNA samples on the chip, a mouse miRNA chip (v.12.0, Agilent Company) was applied. GeneSpring GX v.12.1 software package (Agilent Technologies, USA) was used for data processing. Differentially expressed miRNAs that were statistically significant between the sham and tumor groups were detected via Volcano Plot filtering. Furthermore, differentially expressed miRNAs among two different samples were detected via Fold Change filtering.

### Bioinformatics analysis

The genes that are potentially targeted by miR-135-5p were indicated by miRanda (http://www.mirdb.org/), TargetScan (http://www.targetscan.org/), PicTar (https://pictar.mdc-berlin.de/), RNA22 (https://cm.jefferson.edu/rna22/), PITA (https://genie.weizmann.ac.il/), and RNA-seq results, and the combination of target genes of these five groups was obtained. Cytoscape software (v.3.6.1) was used to create and analyze a miRNA-hub gene network to explore the relation between potential target genes and candidate DE-miRNAs. The DAVID bioinformatics program (https://david.ncifcrf.gov/tools.jsp) was utilized to conduct analysis of the Kyoto Encyclopedia of Genes and Genomes (KEGG) pathways to predict the influences of the downregulated mRNAs.

### Transfection reporter assay

The bioinformatics prediction website (http://www.microRNA.org) was utilized to estimate the target relationship between the JAK2 gene and miR-135-5p and dual luciferase reporter gene assay was utilized for verification. The segment of the JAK2-3’-untranslated region (3’UTR) gene was produced artificially and embedded inside the pMIR-reporter via HindIII and SpeI endonuclease sites. The mutation locations of the complementary sequence of the respective seed sequence were created in wild-type (WT) JAK2. Following treatment with restricted endonuclease, T4 DNA ligase was utilized to incorporate the target segment inside the plasmid pMIR-reporter. The luciferase reporter plasmids which were accurately sequenced, containing WT and mutated (MUT), were cotransfected with miR-135-5p mimics into cultured primary astrocytes. These cells were collected and lysed after 48 h of transfection, and a luciferase assay kit (GM-040501A; Genomeditech Co., Shanghai, China) was used to assess luciferase activity.

### Flow cytometer

To evaluate apoptosis quantitatively, cells were first washed with a solution of cold PBS and then binding buffer was used for resuspension. Following staining of the samples with 10 ml PI and 5 ml annexin V-FITC, an EpicsAltra flow cytometer (Beckman Coulter, CA, USA) was utilized to categorize the cells according to their staining pattern. Apoptosis was indicated when negative PI staining and positive annexin V-FITC staining was observed. Necrosis was indicated when both staining types were positive.

### 5-Ethynyl-2’-deoxyuridine (EdU) analysis

Following management of the cells in accordance with the time and treatment conditions of each group, EdU (Sigma-Aldrich) medium, at a concentration of 50 μmol/L, was added to every dish. An incubator set at 37□ with 5% CO_2_ was used to culture the cells for 2 h followed by rinsing with PBS two times. Then the cells were fixated with 4% paraformaldehyde, replenished with 0.2% glycine, and with PBS rinsed for 5 min in total. The membrane was parted by 0.5% Triton-100 for a total of 10 minutes, followed by rinsing with PBS and replenishment of 100 μl solution of Apollo staining reaction to every well. The cells were incubated, in an environment at room temperature, in a shaker for the duration of 30 minutes in the dark, and after rinsing with 0.5% Triton-100, they were washed with PBS and 100 μl of methanol. Subsequently, staining of the cells was achieved with Hoechst 33258. The rate of EdU incorporation was defined as the fraction of EdU positive cells (red color) i the total number of Hoechst 33258-positive cells (blue color).

### Fluorescence in situ hybridization

Locked nucleic acid (LNA) probes, which corresponded to miR-135-5p were marked with 5’ and 3’-digoxigenin (Exiqon, Woburn, MA, USA). The spinal cord tissue of mice was used for fluorescence in situ hybridization (FISH). After taking this section out, the gene break probe was removed. Then, it was transferred to the instrument for hybridization for 10 min at 75°Cand 42°C overnight. The next day it was removed and rinsed for 5 minutes in an environment at room temperature and for 3 minutes at 72°C. Then, it was dehydrated and replenished drop-by-drop with DAPI to coat the slide. To read and take images, a fluorescence microscope (Olympus IX-81; Olympus, Tokyo, Japan) was used. Then the miR-135-5p cells were analyzed in three separate fields of a single sample that were of high-power and considered to be representative.

### Western blotting

Cells, which were observed to be in the phase of logarithmic growth were collected and seeded in a 60 mm petri dish. Following cell treatment during 24 h, the cells of each of the groups were retrieved to undergo lysis with RIPA. The BCA method was employed to disclose the protein concentration of the cells of each group and 6%-12% SDS-PAGE electrophoresis was performed with an amount of protein of 20 μg/well. The proteins that were isolated by electrophoresis, were subsequently moved to the PVDF membranes. Following incubation of the first antibody, we incubated the second antibody anti-rabbit immunoglobulin G (diluted 1: 5000; Abcam; ab99697; USA) for 1 h in an environment at room temperature and used ECL reagent (Thermo Fisher Scientific, Inc.) for development. To obtain the optical density, each protein band’s gray value was measured by Image J software (National Institutes of Health).

The primary antibody data was as follows: anti-JAK2 antibody (diluted 1: 5000; Abcam; ab108596; USA), anti-p-JAK2 antibody (diluted 1: 1000; Abcam; ab32101; USA), anti-STAT3 antibody (diluted 1: 1000; Abcam; ab68153; USA), anti-p-STAT3 antibody (diluted 1: 2000; Abcam; ab76315; USA), anti-GFAP (diluted 1: 10000; Abcam; ab7260; USA), anti-TNF-α antibody (diluted 1: 800; Abcam; ab66579; USA), anti-IL-1β antibody (diluted 1:1000; Abcam; ab9722; USA), anti-IL-6 antibody (diluted 1:1000, Abcam; ab6672; USA), anti-β-actin (diluted 1:5000; Abcam; ab8227; USA).

### Cellular immunofluorescence and TUNEL staining of tissue sections

The mice under deep anesthesia were transcardially perfused with 0.9% saline first and then 4% formaldehyde. Segments of spinal cord at L4 to L5 were taken out and fixed in 4% formaldehyde overnight. A vibratome (Vibratome 1000 Plus; Vibratome Co, St. Louis, MO) was used to cut sections of 30 μm. For staining with immunofluorescence, free-floating portions were first blocked with Tris buffered saline (TBS) supplemented by 5% goat serum or 5% donkey serum, based on the host species of the secondary antibodies, for 2 h in total and then incubated overnight at 4°C in primary antibody. Next, the portions were washed in PBST (Phosphate Buffered Saline with Tween 20; thrice for 5 minutes) and incubated at room temperature in the secondary antibody for a total of 2 h and washed. The dilution of primary antibodies used included p-JAK2 (diluted 1: 1000; Abcam; ab32101; USA) and GFAP (diluted 1: 50; Abcam; ab4648; USA). Then, the sections were placed on slides and DAPI (Abcam, USA) was used for incubation. Images were obtained a kit under a Zeiss LSM780 confocal microscope (CarlZeiss, Oberkochen, Germany). To detect apoptotic activity, a kit (Promega, Fitchburg, WI, USA) was used to perform TUNEL staining in accordance with manufacturer’s instructions.

### Statistical analysis

SPSS 19.0 software (SPSS Inc., Chicago, Illinois, USA) was used to analyze all the data and the experiments were replicated thrice. The results were stated as mean ± SD. Independent sample t test was conducted to compare the two groups and one-way analysis of variance (ANOVA) to compare between a multitude of groups. P<0.05 was defined as statistically significant.

## Results

### Pain hypersensitivity in BCP mice

NCTC 2472 sarcoma cells were embedded inside the intrafemur to construct a BCP mouse model. Adjustments in nociceptive behaviors were estimated by establishing the PWMT (Fig. 1a) and NSF (Fig. 1b) on days 0 (as baseline), 4, 7, 10, 14, 21, and 28 post surgery. The PWMT and NSF at baseline did not significantly differ in the tumor and sham group (P>0.05, n=8, respectively). The recovery of nociceptive behaviors observed on day 7 and onwards in the sham group indicate that the acute hyperalgesia may have been related to the surgery. The results in the tumor group show that the PWMT decreased from day 10 (P<0.001 and n=8, respectively), while the NSF increased continuously for 28 days (P<0.001 and n=8, respectively). Both the PWMT and NSF indicated statistically significant differences in comparison to the sham group on days 10, 14, 21, and 28 post surgery (P<0.001 and n=8, respectively). Femur slices stained with hematoxylin and eosin showed infiltrated tumor cells inside the marrow cavity, interrupted bone trabeculae, and damaged bone cortex in BCP mice on the 14th day post-surgery (Fig. 1c).

**Fig. 1.**
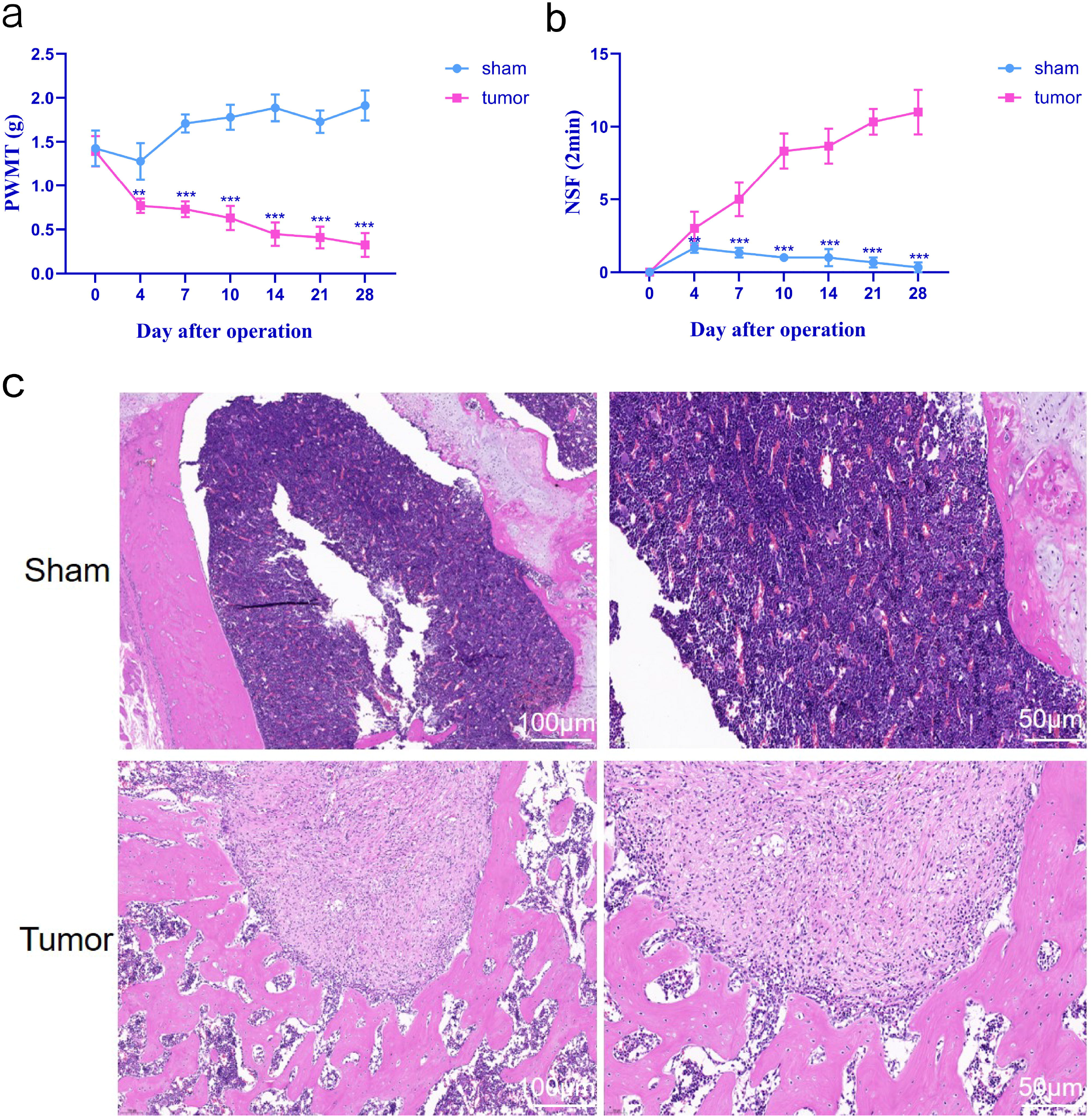
Mechanical hypersensitivity in the homolateral hind paw resulting from the intrafemur implantation of NCTC 2476 cells. (a and b) The paw withdrawal mechanical threshold (PWMT) and number of spontaneous flinches (NSF) were calculated on days 0, 4, 7, 10, 14, 21, and 28 post surgery in the tumor and sham groups. The results of the One-way ANOVA with post hoc test, **P < 0.01, ***P < 0.001 in comparison to the sham group at every point; n = 8 per group. (c) The photomicrographs of the femur’s marrow cavity, of the sham and tumor group, after staining with hematoxylin and eosin on day 14 post surgery. An obvious degeneration of bone and bone marrow replaced by sarcomatous cells can be seen in the tumor group. Data are stated as mean ± SD.

### Decreased expression of miR-135-5p in BCP mice

To determine the function of miRNA in BCP, we first used miRNA microarray to analyze the expression profiles of miRNA on the spinal cord tissues from tumor group vs. sham group (Fig. 2a). Using these significantly dysregulated miRNAs to perform unsupervised clustering analysis made it possible to distinguish BCP mice from controls (Fig. 2b). We first tested these potential miRNAs with a separate cohort of 5 sham controls and 5 BCP mice. To perform further analysis, only the miRNAs with a p value <0.01 and a mean fold change of >5 or <0.2 were selected. Based on the abovementioned criteria, it was found that miR-135-5p, miR-19-5p, miR-181c-3p, and miR-770-5p were significantly dysregulated (Table 1). Then, we used qRT-PCR to further evaluate these four miRNAs in an additional independent cohort consisting of 12 BCP mice and 12 sham controls. It was found that among the four miRNAs, miR-135-5p was significantly downregulated in BCP mice in comparison to the sham controls (Table 1, Fig. 2c), which was confirmed again by FISH (Fig. 2d). The expression level of miR-135-5p were further analyzed in different cells in BCP model vs. normal control. Only in astrocytes, miR-135-5p was significantly differentially expressed (Fig. 2e). Fig. 2f showed the co-localization of miR-135-5p with GFAP, which is the cell marker of astrocyte. These outcomes suggest that miR-135-5p may have disease-specific roles in BCP.

**Fig. 2.**
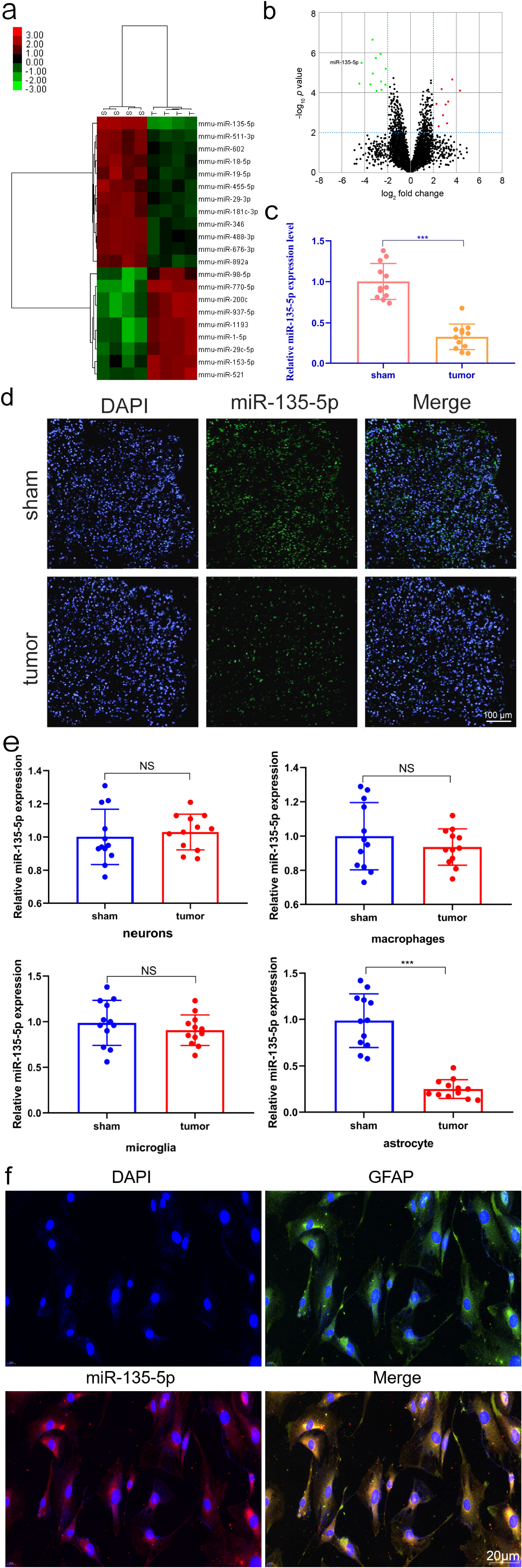
Detection of differentially expressed miRNAs in spine cord tissues of BCP mice. (a) Heatmap was produced by unsupervised clustering analysis with 21 substantially dysregulated miRNAs in on spinal cord tissues from tumor group. Hierarchical clustering was conducted with mean linkage and uncentered correlation. mmu, mouse. (b) Volcano plot illustrates the statistical and biological significance of the expression levels of differential miRNA between tumor and sham groups. Green dots mark the downregulated (left side) and red dots mark upregulated (right side) miRNAs. miR-135-5p is indicated. (c) The expression level of miR-135-5p in 12 BCP mice and 12 sham controls (in the validation and training sets), detected by qRT-PCR assay (***P < 0.001). (d) The results were further verified via in situ hybridization. (e) Only in astrocytes, miR-135-5p was significantly differentially expressed. (f) The co-localization of miR-135-5p with GFAP, which is the cell marker of astrocyte. All data are expressed as mean ± SD.

**Table 1.**
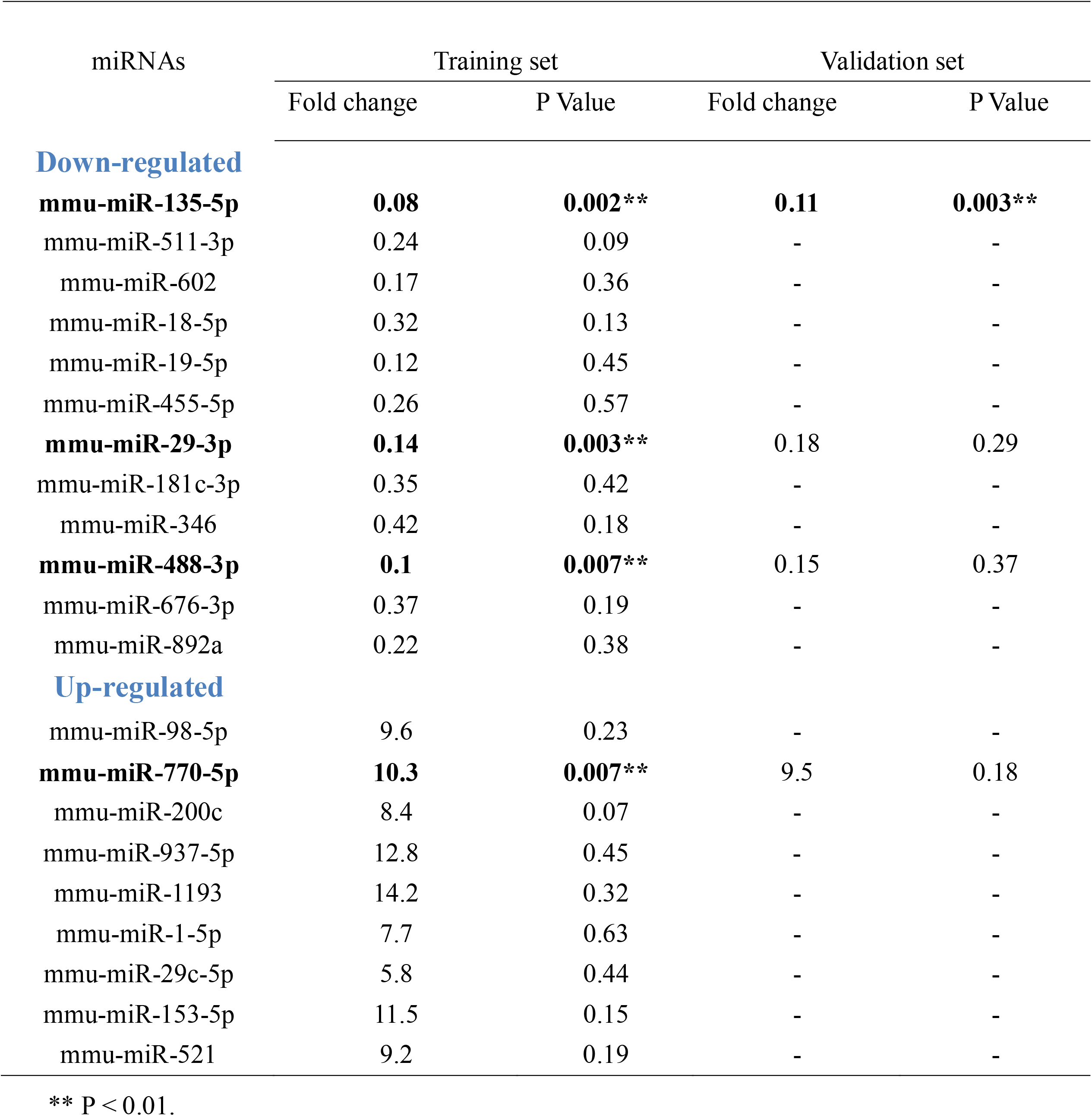
Differentially expressed miRNAs in both the training set and the validation set

### The effect of miR-135-5p silencing or overexpression on astrocytes phenotype

To gain more understanding of the functional role that miR-135-5p plays in BCP pathogenesis, we transfected miR-135-5p inhibitor or mimics inside primary astrocytes which were cultured. The efficiency of transfection was identified by Cy3-labeled miRNA (Fig. 3a). EdU assay was conducted to evaluate the functions of miR-135-5p in the proliferation of astrocytes. The results of our analysis have shown that miR-135-5p inhibitor predominantly stimulated the proliferation of astrocytes compared to its corresponding inhibitor control, whereas miR-135-5p overexpression markedly decreased the proliferation of astrocytes (Fig. 3b). In terms of astrocytes apoptosis, it was found that miR-135-3p mimics transfection significantly increase the amounts of astrocytes apoptosis (Fig. 3c). The effects were further confirmed by immunofluorescence (Fig. 3d). Moreover, by performing gain-of-function and loss-of-function studies, we continued analyzing the impact of miR-135-5p expression on catabolic/anabolic markers. The findings showed that in astrocytes transfected with miR-135-5p inhibitor, expression of the cytokines (TNF-α, IL-1β, and IL-6) was increased. On the contrary, overexpression of miR-135-5p significantly diminished the amount of TNF-α, IL-1β, and IL-6 (Fig. 3e). Overall, the results suggest that downregulating miR-135-5p induces astrocytes proliferation and inflammation factors expression.

**Fig. 3.**
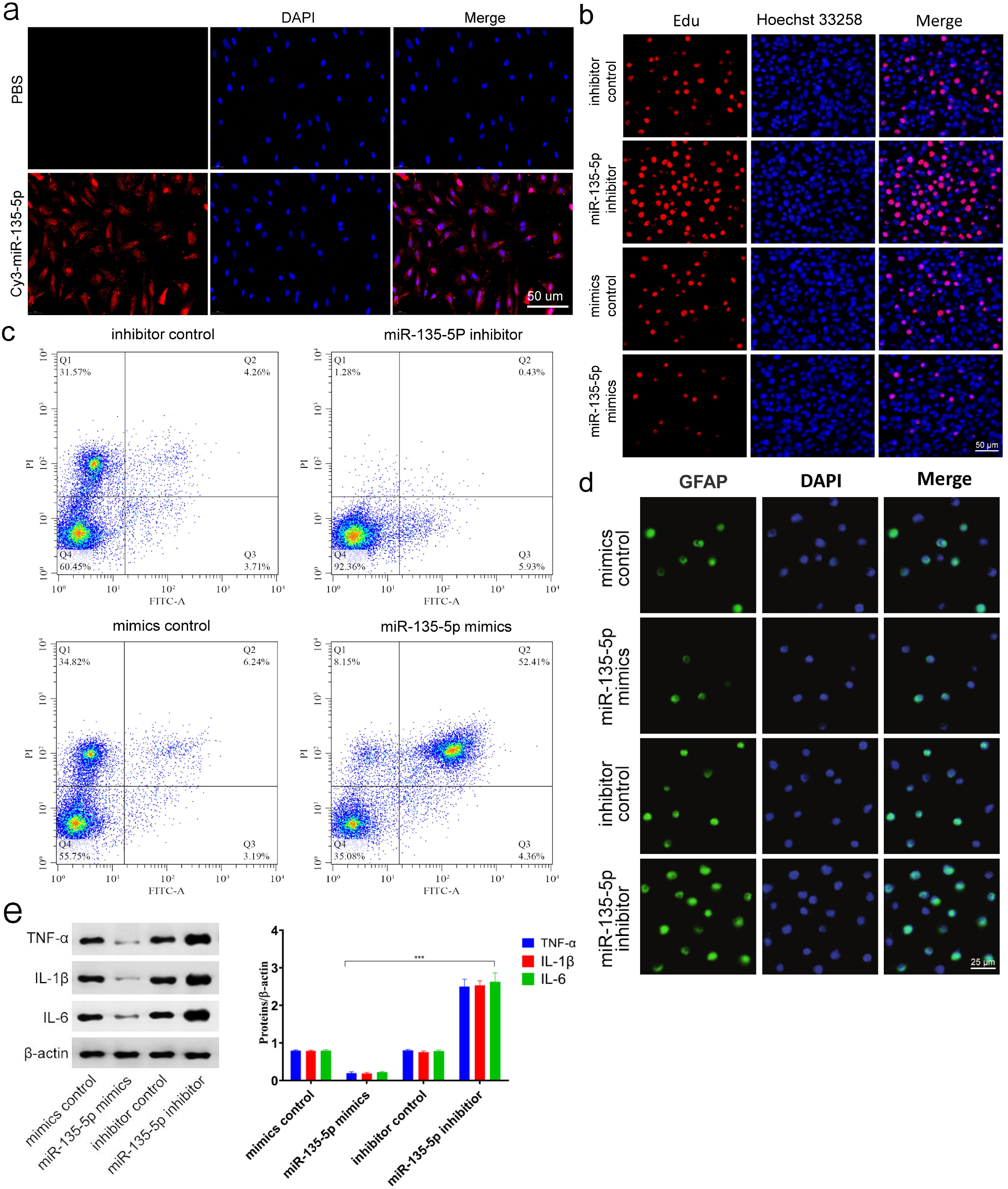
In vitro functional analysis of miR-135-5p. (a) miR-135-5p transfected with cultured primary astrocytes as established by Cy3 label. (b) Analysis of cell proliferation in cultured primary astrocytes transfected with miR-135-5p mimics or inhibitor by EdU assay, with n = 3 replicates per group. (c) Analysis of astrocytes apoptosis was performed by flow cytometer, with n = 3 replicates per group. (d) The representative GFAP were detected by the immunofluorescence. (e) Western blot was used to identify TNF-α, IL-1β, and IL-6 expression levels.

### Identification of JAK2 as a target gene of miR-135-5p

Through searching for possible targets of miR-135-5p, we gathered every predicted gene to conduct Venn analysis (Fig. 4a). Furthermore, a miRNA–mRNA network was constructed by Cytoscape software (Fig. 4b). The miRNA targets predicted by the bioinformatic software and database showed that there is a direct binding between miR-135-5p and JAK2 (Fig. 4c). Analysis of the luciferase reporter assay was conducted to additionally validate the functional relation between miR-135-5p and JAK2. The relative luciferase reporter activity was significantly lower in co-transfected JAK2 WT with miR-135-5p mimics in astrocytes than that of co-transfected JAK2 MUT with miR-135-5p mimics cells (Fig. 4d). This result was also confirmed by the expression of protein in astrocytes (Fig. 4e).

**Fig. 4.**
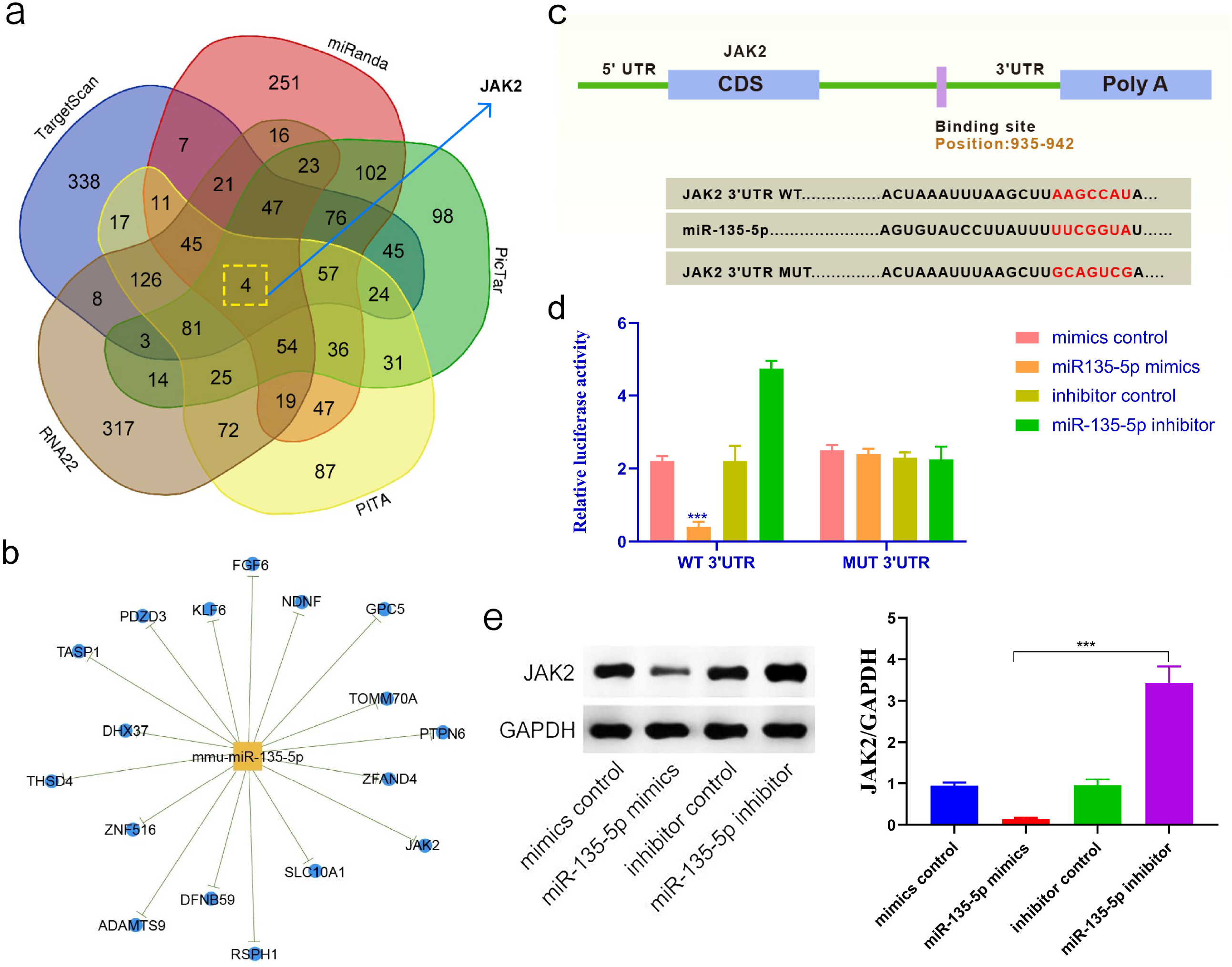
Identifying JAK2 as a miR-135-5p target. (a) Venn diagram shows the computed prediction of miR-135-5p targeting JAK2 via various algorithms. (b) Cytoscape was utilized to establish miR-135-5P’s target. (c) Sequence arrangement of a predicted miR-135-5P binding site among the 3’UTR of JAK2 mRNA indicates a great level of complementarity and sequence preservation with miR-135-5P. (d) The wild- or mutant-type JAK2 3’UTR reporter plasmid was either co-transfected with miR-135-5P mimics, or inhibitor into primary astrocytes, which were cultured. Following transfection, the luciferase activity was calculated, with n = 3 replicates per group. ***P < 0.001 analysis conducted with the one-way ANOVA and post hoc test. (e) Western blot was performed to identify the expression levels of JAK2 in astrocytes. Data are expressed as mean ± SD.

### MiR-135-5p regulates BCP by modulating JAK2/STAT3 signaling pathway

As Fig. 5a shows, the JAK2/STAT3 signaling pathway was enhanced significantly in KEGG pathways. The findings that silence of miR-135-5p promotes BCP mediated by JAK2 motivated us to analyze the potential relation between miR-135-5p and the JAK2/STAT3 pathway. The astrocytes were first transfected with either miR-135-5p mimics, miR-135-5p inhibitor, or their corresponding negative control, respectively. The outcomes validated that overexpressed miR-135-5p led to a reduction in JAK2 and STAT3 expression; whereas, silenced miR-135-5p led to an increase in JAK2 and STAT3 expression (Fig. 5b). Moreover, the level of expression s of GFAP, TNF-α and IL-1β were significantly decreased in astrocytes that steadily overexpress miR-135-5p (Fig. 5b). While, the level of expression s of GFAP, TNF-α and IL-1β were upregulated in astrocytes transfected with miR-135-5p inhibitor (Fig. 5b). In addition, to establish the effects of JAK2 on miR-135-5p in terms of GFAP, TNF-α and IL-1β expression, a rescue test was performed by first overexpressing just miR-135-5p and then overexpressing miR-135-5p as well as JAK2. The overexpression of JAK2 successfully countered the effects of overexpressed miR-135-5p (Fig. 5c).

**Fig. 5.**
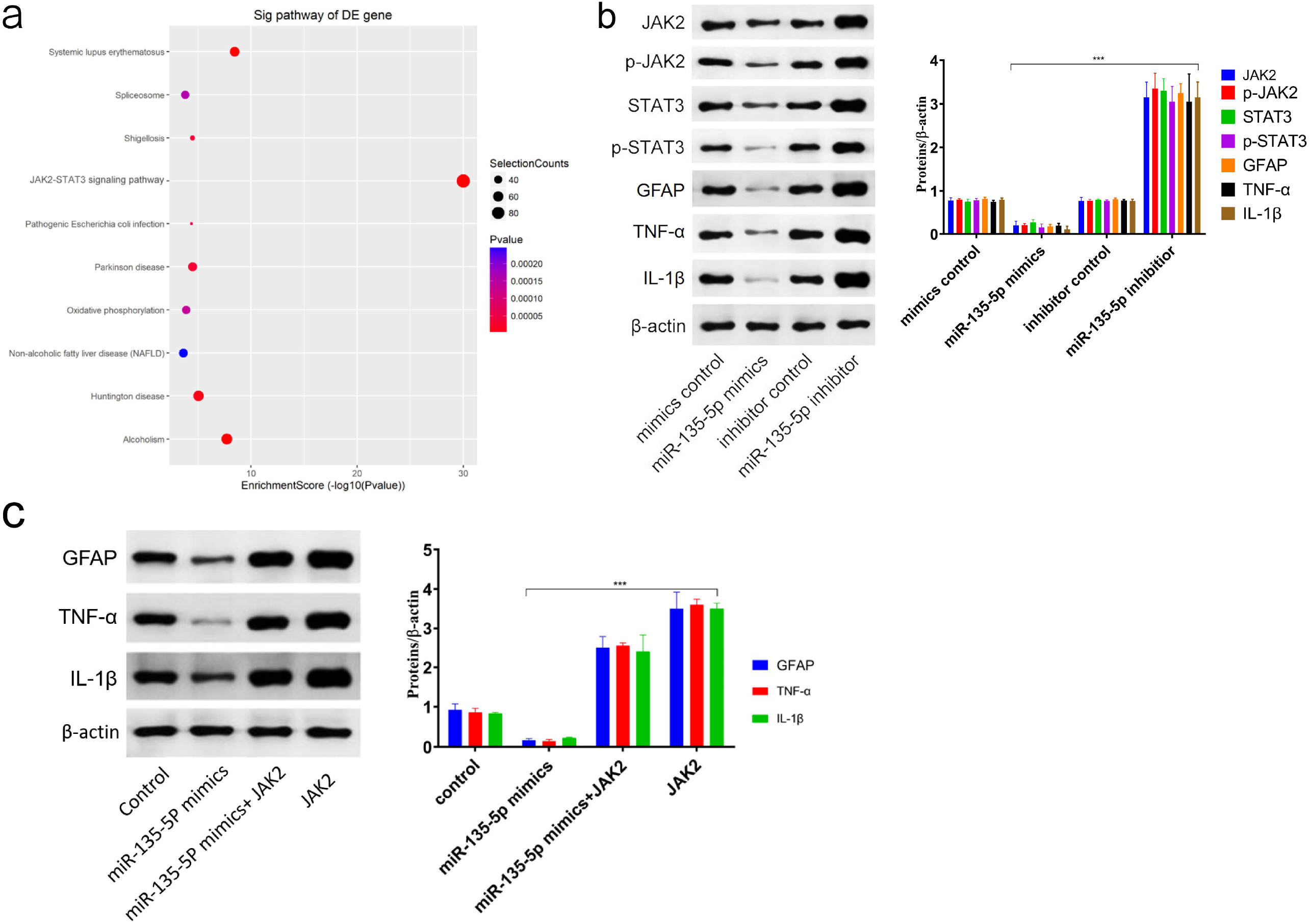
The effect of miR-135-5p on the JAK2/STAT3 signaling pathway. (a) The KEGG analysis showed that the JAK2/STAT3 pathway was enhanced in the BCP model. (b) Cultured primary astrocytes, which were first transfected with miR-135-5p mimics, miR-135-5p inhibitor, or their negative controls, followed by measurements of the levels of JAK2, p-JAK2, STAT3, p-STAT3, GFAP, TNF-α and IL-1β with Western blotting. (c) The rescue experiments were performed in cultured primary astrocytes to verify the correlation between miR-135-5p and JAK2. Suppression of the expression levels of GFAP, TNF-α and IL-1β by miR-135-5p mimics was recovered by restoring JAK2 expression levels.

### Intrathecal administration of miR-135-5p agonist or antagonist in a BCP mouse model

We investigated the potential therapeutic capacity of miR-135-5p in BCP and to clarify which molecular processes participate herein, a BCP model was first constructed in WT mice and then intrathecal injections of miR-135-5p agonist or antagonist and negative control were given at L4–L5 lumbar vertebrae levels every 2 days (three injections). Local delivery of miR-135-5p agonist remarkably increased PWMT and decreased NSF in the BCP group (Fig. 6a and 6b), suggesting that overexpression of miR-135-5p had a preventive effect against BCP. In contrast, mice which were treated with miR-135-5p antagonist activated JAK2/STAT3 signaling pathway. Moreover, GFAP, TNF-α, IL-1β, and IL-6 expression was significantly increased by miR-135-5p antagonist (Fig. 6c). Opposing outcomes were found in mice, which were treated with miR-135-5p agonist (Fig. 6c). These findings were confirmed by immunofluorescence (Fig. 6d). TUNEL staining indicated evidently increased apoptosis of astrocytes in mice treated with miR-135-5p agonist (Fig. 6e). Overall, the outcomes reveal that overexpression of miR-135-5p significantly attenuated BCP, highlighting miR-135-5p as a target with therapeutic potential in BCP.

**Fig. 6.**
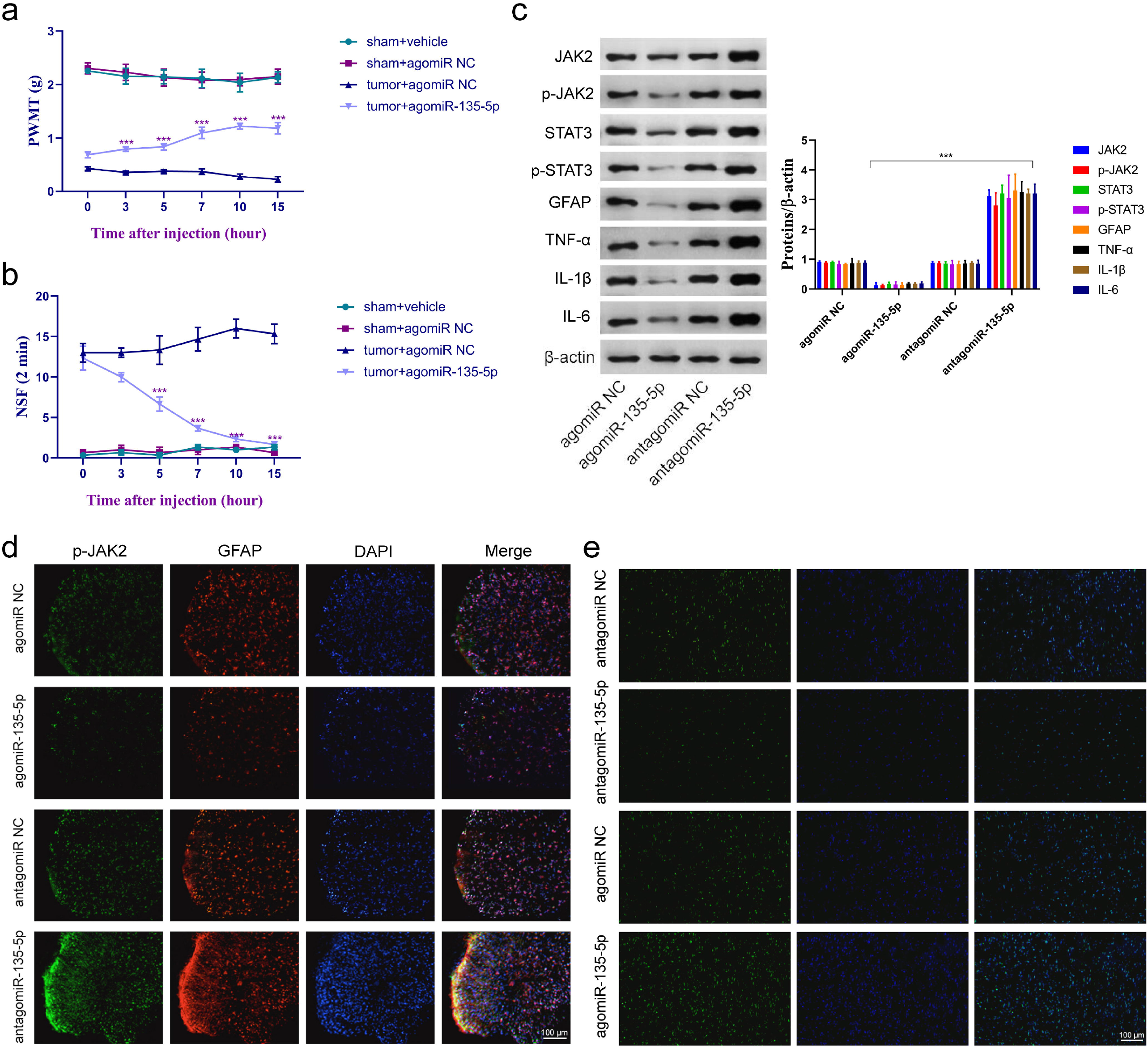
The local deliverance of miR-135-5p agonist decreased the development of BCP. (a and b) Paw withdrawal mechanical threshold (PWMT) and the number of spontaneous flinches (NSF) were measured prior to administration (0 h) and at 3, 5, 7, 10 and 15 h following administration. Analysis performed by One-way ANOVA with post hoc test, ***P < 0.001 in comparison to the mice of the tumor group which were treated with negative control at every point, with n = 12 per group. (c) The levels of JAK2, p-JAK2, STAT3, p-STAT3, GFAP, TNF-α, IL-1β and IL-6 were calculated by Western blotting. (d) Immunostaining of p-JAK2 and GFAP in BCP model treated with miR-135-5p agonist or antagonist. (e) TUNEL staining demonstrated the effect of miR-135-5p overexpression on apoptosis in vivo. Data are expressed as mean ± SD.

## Discussion

It has been demonstrated that a wide range of miRNAs are significant posttranscriptional regulators with vital roles to play in human diseases. Their potential effects on BCP has been the focus of multiple studies [5,29,6,30]. In this study, we studied the role of miR-135-5p in BCP. With a reliable mouse BCP model (Fig 1), we have shown that miR-135-5p expression was reduced significantly in BCP mice. Overexpression of miR-135-5p by treating with a miR-135-5p agonist reduced activation of astrocytes and expression of inflammation factors. Conversely, downregulation of miR-135-5p by its antagonist induced astrocyte-mediated neuroinflammation. Lastly, it was demonstrated that JAK2 was targeted in a direct manner by miR-135-5p. The findings have revealed that miR-135-5p plays a significant part in astrocyte-mediated neuroinflammation, and changes in the expression of miR-135-5p contribute to BCP.

Microarray detection of astrocytes in BCP was performed to investigate the difference in miRNA expression profile between that of BCP and normal spinal cord tissues (Fig 2a and 2b). MiR-135-5p is highly related to the manifestation and progression of numerous diseases [10,31,11,32-35]. In the present study, the impact of miR-135-5p on the manifestation and progression of BCP was assessed. To additionally explore the variance of miR-135-5p in BCP, the expression level of miR-135-5p was detected with qRT-PCR, and the results showed that downregulation of miR-135-5p in BCP existed (Fig 2c). These results were also verified by FISH maps (Fig 2d). Furthermore, the expression level of miR-135-5p were analyzed in different cells in BCP model vs. normal control. Only in astrocytes, miR-135-5p was significantly differentially expressed (Fig 2e). To elucidate the effect of miR-135-5p on astrocytes, the effect of miR-135-5p overexpression and suppression on the astrocyte phenotype was investigated through regulation of miR-135-5p levels. The outcomes showed that suppression of miR-135-5p is related to enhanced proliferation of astrocytes, suppression of astrocytes apoptosis, and upregulation of inflammatory cytokines (Fig 3). These changes in phenotype are considered as vital biological processes in BCP. Thus, miR-135-5p plays a part in the pathogenesis of BCP through astrocytes.

It is known that miRNAs can take part in networks of gene expression by interacting directly with the 3′UTR of targeted mRNAs [36]. Therefore, we selected the candidate direct gene targets of miR-135-5p in databases, which have previously been reported as participants in the pathogenesis of BCP (Fig 4a). To further investigate in what way miR-135-5p regulates BCP, a diagram depicting the miRNA-mRNA network was produced (Fig 4b). JAK2 was detected as the most promising target according to the search results and comparison of the algorithms of sequence complementarity (Fig. 4c). In the astrocytes, the luciferase enzyme activity was reduced significantly following transfection with miR-135-5p mimics in comparison to the cells that were transfected with the mimic control (Fig. 4d). After transfection with the miR-135-5p inhibitor, the enzymatic activity was enhanced notably in comparison to the cells that were transfected with the inhibitor control (Fig. 4d). These findings show that miR-135-5p can interact directly with the 3′UTR of JAK2 mRNA, which adequately suppressed the subsequent translational activities from the chimeric transcript. Western blotting results verified that JAK2 expression in astrocytes was decreased by the miR-135-5p mimics, an oligonucleotide sequence constructed to function similarly to the primary intact structure of miR-135-5p (Fig. 4e). These results have shown that JAK2 serves as a downstream mediator, in the transduction of signals from miR-135-5p, in the pathogenesis of BCP.

Multiple studies have suggested that the JAK2/STAT3 signaling pathway is related to gliocyte proliferation during pain states [24,37,38]. In Tsuda’s study, it was clearly shown that the JAK2/STAT3 pathway in astrocytes is vital for the management of astrocyte proliferation following an injury of the peripheral nerve [14]. JAK2 belongs to a novel family of cytoplasmic tyrosine kinases which has been involved in cytokine receptors and catalyzation of downstream STAT3 activity through phosphorylation [39]. In addition, in reaction to various cytokines, such as TNF-α, IL-6, and IL-1β, the STAT3 pathway becomes activated during the inflammatory response both in vitro and in vivo [40-42]. In our study, KEGG pathway analysis showed that the JAK2/STAT3 signaling pathway was markedly enriched in astrocytes of BCP (Fig. 5a). Moreover, we found that miR-135-5p mimics significantly decreased phosphorylation of JAK2 and STAT3, in addition to the expression of GFAP, IL-1β, and TNF-α (Fig. 5b). However, this effect of miR-135-5p mimics was abrogated by JAK2 treatment (Fig. 5c). Therefore, through functional studies performed in vitro, we have verified that miR-135-5p suppresses the JAK2/STAT3 signaling pathway in BCP.

Our primary hypothesis derived from our results is that dysregulated levels of miR-135-5p could serve as an effective target in the BCP treatment strategy. The significance of this promising hypothesis clinically could be verified further by experiments on an animal model. Our results from the behavioral tests indicated that miR-135-5p overexpression can lead to the increment of the threshold of pain in BCP mice as the PWMT increase and reduction in NSF in mice indicate, after treatment with the miR-135-5p agonist. Mice with BCP had a lower PWMT and higher NSF value in comparison with the mice in the sham group (Fig. 6a and 6b). Additionally, the astrocytes and JAK2/STAT3 signaling pathway were activated and TNF-α, IL-1β, and IL-6 expression was increased in BCP mice, while the opposite effect was obtained in BCP+ agomiR-135-5p mice (Fig. 6c). As revealed by histological staining, agomiR-135-5p can effectively suspend the activation of astrocytes and JAK2 signaling pathway after injection into the BCP mice (Fig. 6d and 6e). Thus, it was suggested that miR-135-5p depression is an early event in BCP and might lead to the onset of BCP.

This is a novel study in establishing a correlation between downregulation of miR-135-5p and BCP. Our primary hypothesis deduced from the results of this study is that increasing levels of miR-135-5p as a posttranscriptional regulator could serve as a treatment strategy for BCP. However, one limitation in our study needs to be mentioned. Based on the newest version of the microRNA database miRBase, the miR-135-5p homolog has been detected before in humans. Thus, whether or not miR-135-5p is suitable as a therapeutic target or diagnostic biomarker for BCP recovery in humans needs to be further studied.

Conclusively, this study provides validation and discovery of BCP-specific miRNA transcriptome outlines. We detected downregulation of miR-135-5p in the BCP mice model. Importantly, miR-135-5p inhibits BCP by disrupting astrocyte-mediated neuroinflammation through the JAK2/STAT3 signaling pathway. These outcomes indicate that downregulation of miR-135-5p may have a valuable role in activation of astrocyte-mediated neuroinflammation and could be considered as a potential novel target in BCP therapy.

## Acknowledgements

The authors would like to thank those previous researchers who updated their data onto the databases employed in the present study.

## Authors’ Contributions

ML, XFC, and ZHC designed the experiments. ML and XFC collected samples. ML, XFC and HY conducted the experiments and acquired the data. ML, HY and JLC analysed the data. ML and XFC wrote the manuscript. HY, JLC and CHL revised the manuscript. All authors approved the final version. The corresponding author had full access to all the data in the study and had final responsibility for the decision to submit for publication.

## Availability of data and materials

The datasets generated during and/or analyzed during the current study are available from the corresponding author on reasonable request.

## Ethics approval and consent to participate

All experimental approaches were reviewed and approved by the Institutional Animal Care and Use Committee of the Central Hospital of Wuhan, Tongji Medical College, Huazhong University of Science and Technology.

## Consent for publication

Not applicable.

